# Pervasive introgression of MHC genes in newt hybrid zones

**DOI:** 10.1101/706036

**Authors:** K. Dudek, T. S. Gaczorek, P. Zieliński, W. Babik

## Abstract

Variation in the vertebrate major histocompatibility complex (MHC) genes is crucial for fighting pathogen assault. Because new alleles confer a selective advantage, MHC should readily introgress between species, even under limited hybridization. Using replicated transects through two hybrid zones between strongly reproductively isolated newts, we demonstrated recent and ongoing MHC introgression. Its extent correlated with the age of contact. In the older zone, MHC similarity between species within transects exceeded that of between transects within species, implying pervasive introgression – a massive exchange of MHC genes, not limited to specific variants. In simulations, the observed pattern emerged under the combined action of balancing selection and hybridization, but not when these processes acted separately. Thus, massive introgression at advanced stages of divergence can introduce novel and restore previously lost MHC variation, boosting the adaptive potential of hybridizing taxa. In consequence, MHC genes may be the last to stop introgressing between incipient species.

## Introduction

Speciation is usually a prolonged process, because reproductive isolation between differentiating taxa evolves gradually (Coyne & Orr, 2004). Speciation commonly involves independent evolution in allopatry (Kisel & Barraclough, 2010). However, due to the dynamic nature of species ranges, periods of allopatry are often accompanied by episodes of contact, which set the stage for natural hybridization (Abbott et al., 2013; Harrison & Larson, 2016). If hybridization leads to introgression, some introgressed variants may be beneficial in the recipient species. Introgression of such variants will be favored by natural selection, even if overall introgression is constrained due to low fitness of hybrids and genome-wide barriers to neutral introgression (Barton, 1979). Such adaptive introgression may occur at many loci, perhaps even providing variation that fuels adaptive radiations (Meier et al., 2017), or may be restricted to particular genes or genomic regions (Hedrick, 2013; Huerta-Sánchez et al., 2014). While examples of adaptive introgression have been documented and theory underlying adaptive introgression is available, general properties of the process and the resulting empirical patterns are not sufficiently understood.

Some genes or classes of genes may be particularly prone to adaptive introgression due to the evolutionary processes affecting them. However, paradoxically, convincing demonstration of introgression of these genes may be the most challenging. Genes involved in fighting pathogen assault, which often evolve under various forms of selection that maintain and promote diversity (collectively termed balancing selection, BS), are prime candidates for adaptive introgression (Fijarczyk, Dudek, Niedzicka, & Babik, 2018; Schierup, Vekemans, & Charlesworth, 2000). This is because novel alleles may confer selective advantage and selection would favor their introgression (Phillips et al., 2018). Vertebrate Major Histocompatibility Complex (MHC) genes are a textbook example of long-term BS (Spurgin & Richardson, 2010) driven in a large part by Red Queen dynamics (Ejsmond & Radwan, 2015). Adaptive MHC introgression between hybridizing species has been inferred in several systems (Abi-Rached et al., 2011; Fijarczyk et al., 2018; Grossen, Keller, Biebach, Croll, & Consortium, 2014; Nadachowska-Brzyska, Zielinski, Radwan, & Babik, 2012), with varying strength of supporting evidence. However, studying introgression of targets of balancing selection is challenging because BS itself generates patterns of variation and differentiation resembling those due to introgression, which makes distinguishing between these two processes difficult (Fijarczyk & Babik, 2015). Furthermore, MHC is a dynamically evolving multigene family characterized by frequent duplications and pseudogenizations, resulting in copy number variation and the mixture of functional genes and pseudogenes in the genome (Kelley, Walter, & Trowsdale, 2005). These genomic complexities impede both molecular analyses and realistic modeling and testing of competing evolutionary scenarios.

Hybrid zones – areas where incipient species meet and hybridize – are considered natural laboratories and windows on evolutionary processes, because they allow the outcomes of multiple generations of hybridization under natural conditions to be studied (Harrison & Larson, 2016). Hybrid zones allow direct observation of introgression, comparison between genomic regions, estimation of the strength of genome-wide reproductive isolation, and rigorous testing of adaptive introgression and alternative explanations (Gompert, Mandeville, & Buerkle, 2017; Payseur, 2010). Inferences are especially strong if multiple transects through a hybrid zone are available, which allows for: i) testing the repeatability of patterns between transects, ii) in the case of polymorphic genes, comparing intraspecific divergence between transects with interspecific divergence within transects. The latter constitutes a powerful test for introgression, as increased genetic similarity in contact zones is not expected under any scenario other than introgression (Fraïsse, Belkhir, Welch, & Bierne, 2016; Nadachowska-Brzyska et al., 2012). In the context of testing adaptive introgression of genes under BS, such as the MHC, hybrid zones between strongly reproductively isolated taxa are excellent tools, because theory predicts a pronounced contrast between targets of adaptive introgression and the rest of the genome.

In this study we investigated introgression of MHC genes through hybrid zones between two European salamanders: Carpathian (*Lissotriton montandoni*) and smooth (*L. vulgaris*) newts. These newts hybridize in a narrow contact zone along the Carpathians, but hybridization is currently limited: even at the centre of the hybrid zone, parental genotypes predominate, resulting in a bimodal distribution of genotypes, that indicates strong reproductive isolation (Babik, Szymura, & Rafiński, 2003; Zieliński et al., 2013). However *L. montandoni* and *L. vulgaris* have experienced a long and apparently complex history of genetic exchange (Zieliński, Nadachowska-Brzyska, Dudek, & Babik, 2016), exemplified by a complete replacement of *L. montandoni* mtDNA by that derived from several phylogeographic lineages of *L. vulgaris* (Pabijan, Zielinski, Dudek, Stuglik, & Babik, 2017; Zieliński et al., 2013). Introgression of MHC classes I and II between *L. montandoni* and *L. vulgaris* has previously been inferred using continent-scale comparisons of differentiation and allele sharing between multiple evolutionary lineages within the *L. vulgaris* species complex (Fijarczyk et al., 2018; Nadachowska-Brzyska et al., 2012). However, such large scale analyses provide only limited insight into the temporal and spatial dynamics of introgression. Analyses of MHC introgression in hybrid zones are more informative in this respect, as they allow testing for recent or ongoing introgression and estimation of its strength.

We analyzed zones of hybridization between *L. montandoni* and two evolutionary lineages of *L. vulgaris*: i) *L. montandoni* x *L. v. ampelensis* (the IN zone) and ii) *L. montandoni* x *L. v. vulgaris* (the OUT zone). The two lineages within *L. vulgaris* diverged more than 1 Mya (Pabijan et al., 2017; Zieliński et al., 2016), and each of them has experienced a distinct history of hybridization with *L. montandoni* (Zieliński et al., 2016). In particular, the IN zone appears older (Zieliński et al., 2019). Within each zone we sampled two transects separated by a distance of 100-200 km, which is two orders of magnitude larger than the per generation dispersal distances of these newts. We took advantage of two hierarchically sampled transects through each hybrid zone to compare introgression of MHC genes to that of more than 1100 other protein-coding genes scattered throughout the newt genome.

## Materials and Methods

### Experimental design and sampling

Hierarchical sampling comprised two hybrid zones with two transects within each: i) the IN zone (*L. montandoni* x *L. vulgaris ampelensis*) inside the Carpathian Basin, with the transects Lypcha (L) and Remetea (R) separated by ca 200 km, and ii) the OUT zone (*L. montandoni* x *L. v. vulgaris*) outside the Carpathian Basin, with the transects Suceava (S) and Tazlau (T) separated by ca. 100 km (Fig. 1A). Note that the compound R transect consists of three contact zones, as the two mountain ranges where *L. montandoni* occurs are separated by an upland inhabited by *L. vulgaris*. The description of all transects, including environmental characteristics, is given in Zieliński et al. (2019). We attempted to sample all water bodies where newts were present, with special emphasis on the centre of each zone, where both species occurred in syntopy and individuals of intermediate morphology were found. Each water body was considered a distinct locality and a distinct breeding population, although the distances between localities were often < 1 km, within the newt dispersal ranges. Newts were captured by dip netting during breeding season, tailtip biopsies were taken and preserved in 96% ethanol; animals were released immediately after sampling. DNA was extracted using the Wizard Genomic DNA purification kit (Promega). Data on genome-wide admixture, available for 96% of individuals genotyped in MHC, based on 2849 SNPs (minor allele frequency, MAF ≥ 0.05) in ca. 1100 protein-coding genes are from Zieliński et al. (2019).

**Fig. 1.**
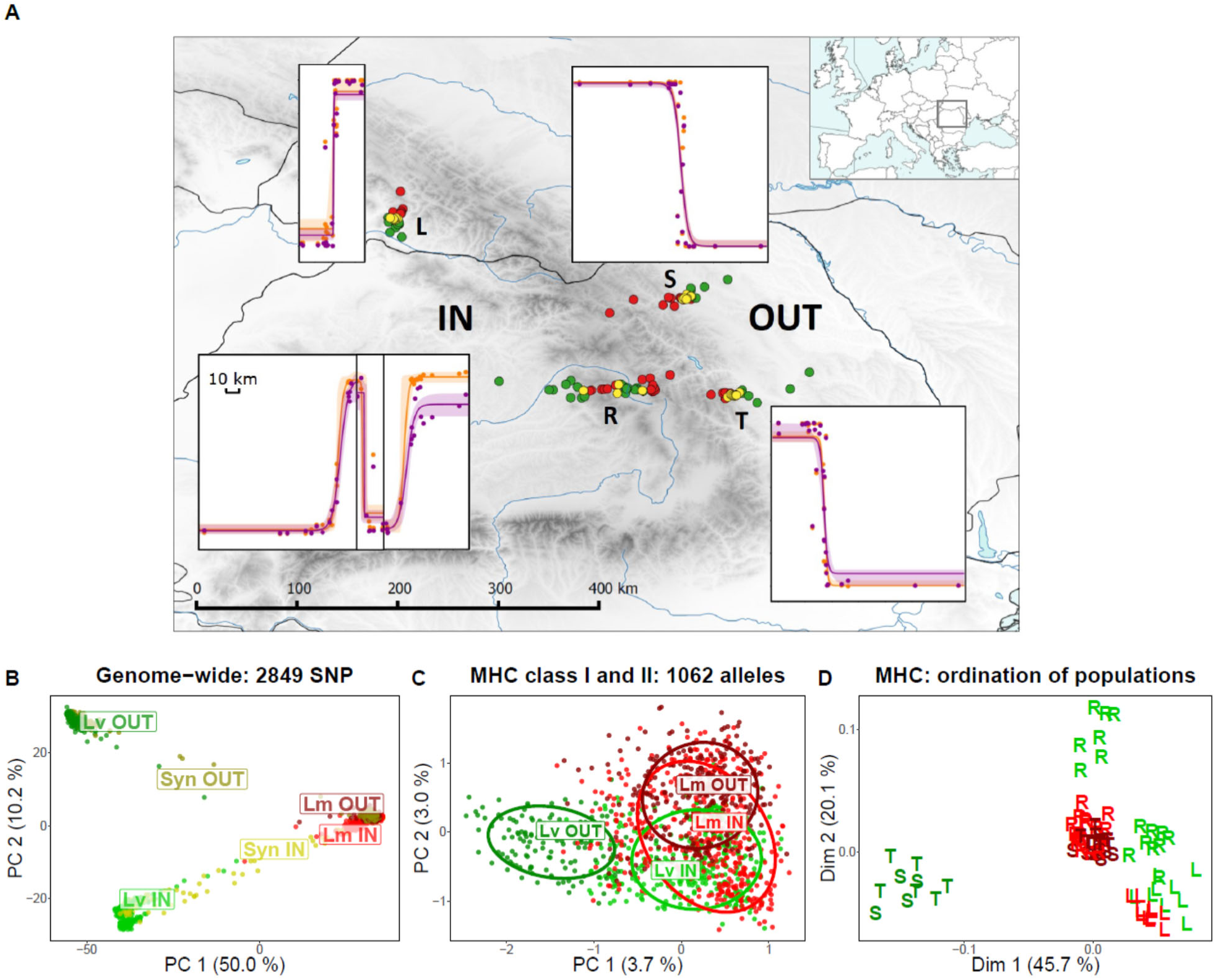
Sampling, clines and differentiation. A) Sampling and clines. Location of the two hybrid zones (IN, OUT), the four transects (L, R, S, T) and localities within transects are shown: green – *L. vulgaris*, red – *L. montandoni*, yellow – syntopic *L. montandoni* & *L. vulgaris*. For each transect genome-wide (orange, mean population Q value from Admixture based on 2849 SNP) and MHC (purple, mean population Q value from Structure based on MHC alleles coded as dominant markers) clines are shown. Note the compound nature of the R transect, which is composed of three contact zones. B) Principal Component (PCA) ordination of individuals based on genome-wide SNP data, light green – *L. vulgaris* inside the Carpathian Basin (IN zone), dark green – *L. vulgaris* outside the Carpathian Basin (OUT zone), light red – *L. montandoni* IN, dark red – *L. montandoni* OUT, yellow – individuals from syntopic populations IN, olive – individuals from syntopic populations OUT; C) PCA based on presence/absence of MHC class I and II alleles (individuals from syntopic populations were excluded); D) Multidimensional scaling ordination of populations based on pairwise F_ST_ in MHC; populations from a single transect are indicated with the same letter.

### MHC genotyping, haplotypes and linkage

The variable 2^nd^ exon of both MHC class I and class II genes was PCR amplified, amplicons were sequenced on the Illumina platform and genotyping was performed using the adjustable clustering method implemented in AmpliSAS (Sebastian, Herdegen, Migalska, & Radwan, 2016). Laboratory and bioinformatics procedures used for MHC class I genotyping were described in detail in Fijarczyk et al. (2018). MHC class II was amplified with primers based on those used in Nadachowska-Brzyska et al. (2012), modified to include additional variation in the regions of primer annealing, as identified by transcriptome resequencing and genome walking. The class II primers (see Supplementary Materials) target a longer fragment than that studied by Nadachowska-Brzyska et al. (2012) (244 vs 204 bp) and we verified, genotyping 9 individuals from that previous study, that the modified primers amplified all alleles previously detected in these individuals. Details of class II genotyping, analyses of diversity and tests of selection are in Supplementary Materials.

Eight MHC class I haplotypes were reported in Fijarczyk et al. (2018). To determine MHC class II haplotypes we used the same individuals: parents and offspring from two families of interspecific *L. montandoni* x *L. v. vulgaris* crosses (11 and 13 F1 offspring). To estimate linkage between the two MHC classes we compared class I and II genotypes in one of the families (parents and 113 offspring) used previously for the construction of a linkage map (Niedzicka, Dudek, Fijarczyk, Zieliński, & Babik, 2017).

### Classical MHC alleles and supertypes

Although the primers were designed from transcriptome sequences, they amplified both highly expressed, putative classical MHC alleles, and alleles expressed at lower level, that may be pseudogenes or nonclassical MHC. Because almost all these alleles segregate as stable haplotypes (Fijarczyk et al., 2018, see also Results), the loci are tightly linked. Therefore all should be useful markers for detecting introgression of the MHC region. Nevertheless, classical genes may undergo different dynamics than nonclassical/nonfunctional genes. It is also possible that alleles within functional classes may be exchangeable and thus affected mainly by drift. Therefore we performed also additional analyses: i) excluding putative nonclassical/nonfunctional MHC class I alleles, and ii) grouping MHC alleles into supertypes (details in Supplementary Materials).

### Statistical analysis

Both MHC class I and class II are duplicated in *Lissotriton* newts and exhibit copy number variation. Assignment of alleles to loci was not possible, therefore we decided to perform statistical analyses on data with each allele encoded as an indicator binary variable (presence-absence). To assess differentiation between species and between transects visually and define main directions of the overall variation we used Principal Component Analysis (PCA) in adegenet. To assess overall MHC differentiation as well as differentiation between species within transects we used multidimensional scaling (MDS) ordination of pairwise F_ST_ distances. To estimate the fraction of variation explained by consecutive levels of hierarchical structuring we employed the AMOVA in Arlequin (Excoffier & Lischer, 2010) with the number of pairwise differences between individuals as the distance measure. Comparison of the width and location of the centre of allele frequency clines is useful to compare introgression between markers. Here we compared MHC clines to the average genome-wide (SNP) clines using the following approach. For each individual the proportion of the genome derived from each species estimated with Admixture (Alexander, Novembre, & Lange, 2009) was taken from Zieliński et al. (2019). The proportion of MHC alleles derived from each species was estimated using Structure (Falush, Stephens, & Pritchard, 2007) with MHC alleles encoded as binary dominant data. For transects L, S and T, two genetic clusters were strongly supported. In the compound transect R (Fig. 1A) three clusters were identified – one for *L. montandoni* and two for *L. vulgaris*, as populations of this species from an upland between two mountain ranges were assigned to a separate cluster. For the purpose of further analyses Q values from the two *L. vulgaris* clusters were summed. Geographic clines were then fitted to the average population Q values for both genome-wide and MHC data using HZAR (Derryberry, Derryberry, Maley, & Brumfield, 2014). For both MHC and SNPs we fitted the same cline model, which estimated the location of cline center (*c*) and width (*w*, 1/maximum slope) as well as the allele frequencies at the ends of the cline. We then used the parameters from the genome-wide cline to test whether clines estimated for MHC data fit the data better, which would indicate differences between clines. The fit was compared using the Akaike information criterion score corrected for small sample size (AICc), with ΔAICc ≥ 5 was considered a significant support for an MHC-specific model.

Isolation by distance between interspecific population pairs within each transect was tested using a matrix permutation procedure (10 000 permutations), in which rows and columns were permuted independently. We tested for an increased allele sharing closer to the centre of the zone in each transect separately. For each species 1-3 (depending on the distribution of localities in the transect) populations at the ends of the transect were classified as allopatric, while the remaining allotopic populations were classified as parapatric. For each species populations within groups were pooled, the fraction of alleles shared between species was calculated for each group and significance of the difference between groups was tested by permuting individuals within species between groups 10 000 times. Because in the OUT zone many syntopic populations were available and most individuals in these populations showed little genome-wide admixture, we also tested for an increased allele sharing in syntopy.

### Simulations

A multiallelic gene under negative frequency dependent selection was simulated according to the infinite allele model in a linear stepping stone (Fig. S1) using Selector (Currat, Gerbault, Di, Nunes, & Sanchez-Mazas, 2015). Although newt MHC is multilocus, its evolutionary dynamics should be reflected adequately by a single, multiallelic locus, because most likely entire haplotypes are subject to selection. To verify this, additional simulations with multilocus MHC haplotypes were performed, as described in Supplementary Methods. An ancestral species split into two equally sized, completely isolated descendant species, which thus initially shared a pool of alleles (Fig. S1I). Following the split, each species colonized a world of demes conforming to a one-dimensional stepping stone model arranged into a horseshoe shape that approximated the two transects of the IN zone, i.e. the distance between transects was 5 times larger than the length of a transect within each species (Fig. S1II). No mutations were allowed, so differentiation between species occurred only via drift, maximizing the retention of ancestral polymorphism. Following prolonged evolution in isolation, secondary contact and hybridization ensued (Fig. S1III). A single deme hybrid zone acted as a partial barrier to gene flow – immigration into the zone was high but emigration from the zone was strongly reduced as in classical tension zone models.

We investigated the effect of the following parameters: strength of selection (s = 0 – 0.3), deme size (N = 100 – 1000), initial number of alleles (na = 15 – 500), migration between demes (Nm = 0.1 – 2.5), strength of barrier to introgression formed by the hybrid zone (emigration from the zone as a fraction of that from demes within species N_h_m = 0.01 – 0.1Nm) and time of hybridization (0.01 – 0.1 of time in isolation). A single value of time in isolation (t = 16 000 generations) was used; because differentiation depends both on time of isolation and population sizes, by varying deme size we also implicitly varied time of isolation. The total number of alleles and the number of alleles within deme stabilized quickly, within ca. 2500 generations, regardless of the deme size.

At the end of each simulation 16 individuals from each deme within each transect were sampled (Fig. S1IV) to calculate: i) fractions of variation (AMOVA) accounted for by the between species and between transect within species levels, ii) the numbers and fractions of alleles shared exclusively (those shared by two focal groups but absent from all other groups) between species within transect and between transects within species. To investigate the temporal and spatial dynamics of introgression we followed the frequency of heterospecific genes through time in a single transect. Details of simulations are in Supplementary Materials.

## Results

### MHC variation and evidence for positive selection

MHC was genotyped in 1672 individuals from 128 localities in four transects (Fig. 1A, Table S1). Genotyping repeatability was 95% (class I) and 98% (class II). Overall variation was high: 920 MHC class I alleles (25 supertypes) in 1679 individuals and 263 class II alleles (22 supertypes) in 1674 individuals (Table S1). The number of MHC alleles per individual ranged from 5 to 25 (class I: 4 – 21, class II: 1 – 7, Fig. S2), indicating extensive copy number variation. Eight haplotypes identified by genotyping two newt families contained 1-3 class II genes, with 5-9 genes per haplotype reported earlier for class I (Fijarczyk et al., 2018).

A strong signal of positive selection was detected in class II (ΔAIC = 74.6 for comparison of PAML models with (M8) and without (M7) positive selection), similar to that reported for class I (Fijarczyk et al., 2018). Only two class II codons were identified as targets of positive selection, but amino acid variation in positions corresponding to the human Antigen Binding Sites was extremely high (Table S2, Fig. S3).

### Comparison between zones and transects: strong MHC introgression in the IN zone

Based on Principal Component (PC) ordination of the genome-wide single nucleotide polymorphism (SNP) data, species were well separated along PC 1, which explained five times more variation than PC 2 (Fig. 1B). In contrast, species were poorly separated on the MHC’s PC1-PC2 plane, even when syntopic populations (defined as localities where both species were identified based on morphology) were excluded from the analyses, and the extent of separation appeared to differ between the IN and OUT zones (Fig. 1C). This interpretation did not change if the MHC classes were analysed separately (Fig. S4), or if MHC supertypes were used instead of alleles. In analyses performed for each zone independently, no separation by species but some separation by transect was observed along PC 1 in the IN zone, while clear separation of species along PC 1 was evident in the OUT zone (Fig. 2A, B). Patterns of allele sharing were consistent with the above picture (Fig. 2C, D). In the IN zone a higher fraction of alleles were shared exclusively (shared by two focal groups but absent from other groups) between species within transect (L: 7.0%, R: 16.2%) than between transects within species (*L. montandoni*: 3.6%, *L. vulgaris*: 2.1%), while in the OUT zone exclusive allele sharing between transects within each species (*L. montandoni*: 17.3%, *L. vulgaris*: 9.7%) was much higher than between species within transect (S: 0.5%, T: 3.2%, Breslow-Day test for homogeneity of odds ratios, IN: *P* = 2.6 × 10^−4^, OUT: *P* = 1.9 × 10^−4^).

**Fig. 2.**
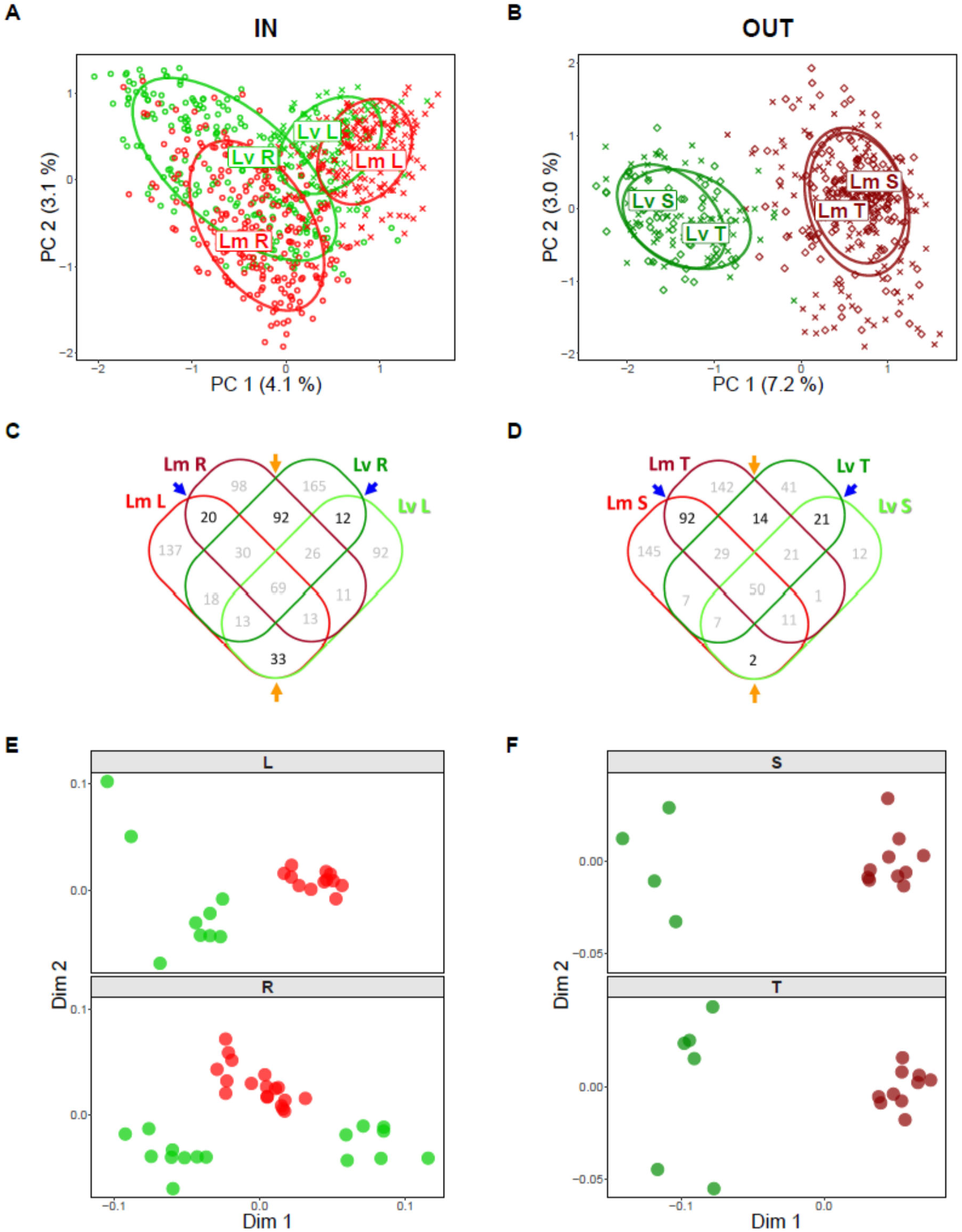
MHC differentiation between species and transects in the IN (A, C, E) and OUT (B, D, F) zones. A) and B) PCA based on presence/absence of MHC class I and II alleles, transects within zones indicated with ellipses and different symbols. C) and D) Venn diagrams showing in black the number of alleles shared between species within transects (orange arrows) and between transects within species (blue arrows). E) and F) Multidimensional scaling ordination of populations within transects based on pairwise F_ST_ in MHC.

Analysis of Molecular Variance (AMOVA) revealed no differentiation between species, but significant differentiation between transects within species in the IN zone. In the OUT zone, the between species component accounted for a substantial fraction of variation (11.6%, Table 1A). The results for supertypes were virtually identical (Table S3). Thus, in the IN zone MHC variation was structured mainly by geography, while in the OUT zone by species (Fig. 1D, 2A, B). In both zones, the genome-wide differentiation between species was much higher than that between transects within species (Table 1A).

**Table 1.**
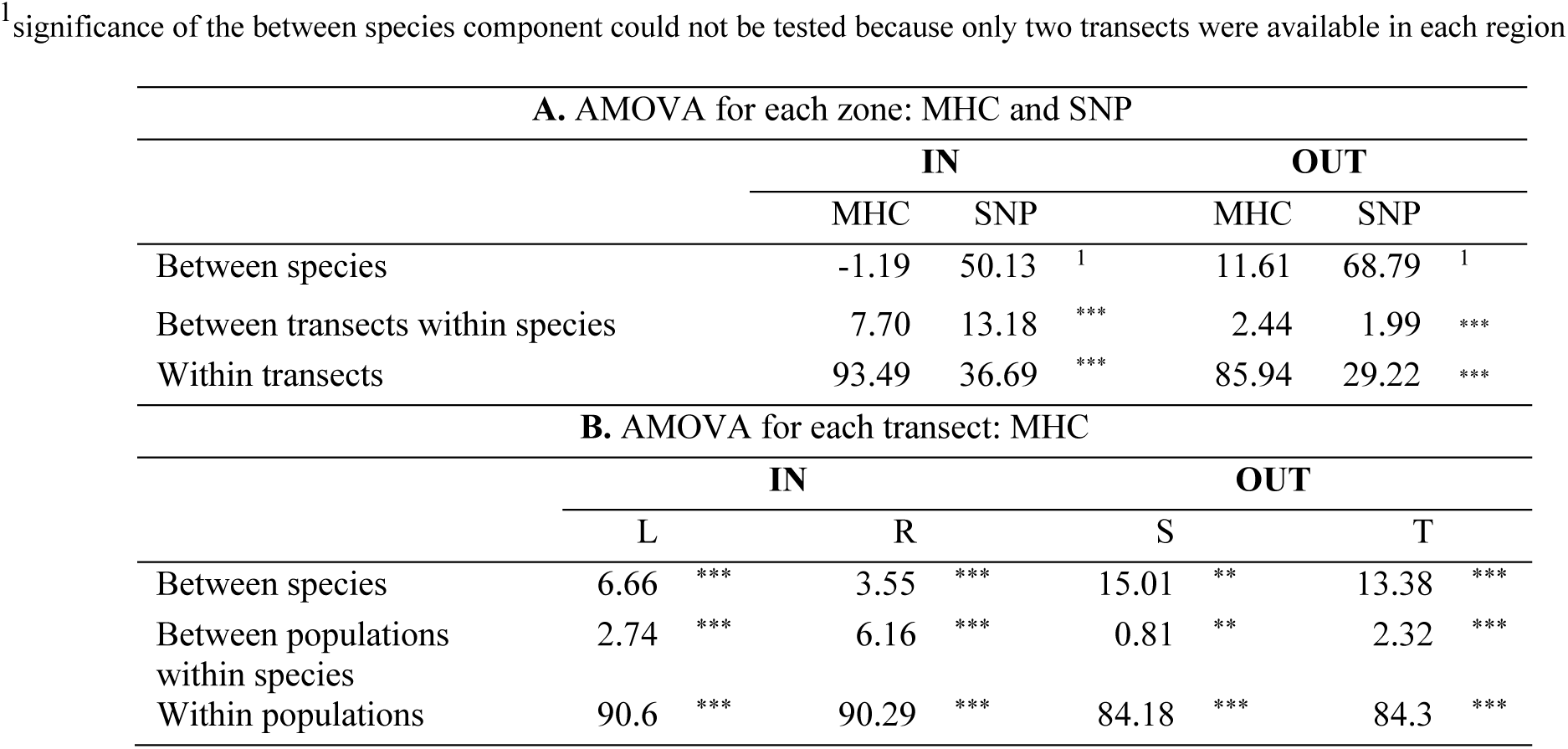
Analysis of Molecular Variance. Percentages of SNP and MHC variation explained by hierarchical levels of population structure are given. Syntopic populations were excluded from the analysis.

### The extent of introgression within transects

Despite strong evidence for introgression in the IN zone, at local scales, i.e. within transects, interspecific differentiation in MHC was visible everywhere, though more so in the OUT zone (Table 1B, S3, Fig. 2E, F). The width or location of MHC clines did not differ from those of genome-wide clines in any transect (Fig. 1A, Table S4). Thus, a sharp transition in MHC, coinciding with the transition in genome-wide ancestry, occurs in each transect. Hence, at local scales, MHC gene flow is currently more restricted between species than between populations within species. However, the magnitudes of MHC and genome-wide differentiation on both sides of the zone are difficult to compare, as Admixture and Structure analyses that we used to estimate genome-wide and MHC ancestry enforce both scales to run from 0 to 1.

If introgression decreased with the distance from the centre of the zone, we would expect significant isolation by distance between heterospecific population pairs within transects. No such pattern was detected in any transect (matrix randomization test, all *P* > 0.05). A complementary, potentially more powerful test, compared the fraction of alleles shared between species closer and further from the centre (Table 2). Importantly, this approach allowed testing for MHC introgression in the OUT zone: although other analyses showed that MHC introgression in the OUT zone was not as strong as the IN zone, they did not provide decisive evidence for or against introgression. In the IN zone, no increased allele sharing was detected closer to the centre of the L transect, but a highly significant increase was found in the R transect. In the OUT zone allele sharing increased towards the centre, providing evidence for MHC introgression (Table 2). Because both species often co-occur in the same ponds in the OUT zone, we were also able to test for increased allele sharing in syntopy, including only newts with little or no genome-wide admixture (< 3%). In the S transect, increased allele sharing was detected only in syntopy, while in the T transect the fraction of shared alleles in allotopy close to the centre was very similar to that in syntopy, and in both transects there was significantly more allele sharing close to the centre than far from the centre (Table 2).

**Table 2.** Allele sharing between species closer and farther from the centre of the hybrid zone. For each comparison gClose is the group closer and gFar is the group farther from the centre (Syntopy < Parapatry < Allopatry). In syntopic populations individuals with > 3% genome-wide admixture were excluded from analysis. One sided *P* values (sharing gClose > sharing gFar, 10 000 permutations of individuals within species between groups) are reported. Lm – *Lissotriton montandoni*, Lv – *L. vulgaris*, Allo – allopatry, Para – parapatry, Syn – syntopy, Tr – transect.

| Zone | Tr | Compared groups |  | Percent alleles shared |  | *P* | Number of MHC alleles |  | Sample size |  |  |  |
| --- | --- | --- | --- | --- | --- | --- | --- | --- | --- | --- | --- | --- |
|  |  | gClose | gFar | gClose | gFar |  | gClose | gFar | Lm gClose | Lm gFar | Lv gClose | Lv gFar |
| IN | L | Para | Allo | 24.2 | 19.2 | 0.80 | 413 | 328 | 145 | 33 | 82 | 33 |
| IN | R | Para | Allo | 39.1 | 18.3 | <0.0001 | 529 | 290 | 222 | 23 | 201 | 25 |
| OUT | S | Para | Allo | 17.3 | 13.7 | 0.23 | 324 | 299 | 102 | 48 | 41 | 19 |
| OUT | S | Syn | Allo | 25.5 | 13.7 | 0.0090 | 271 | 299 | 40 | 48 | 52 | 19 |
| OUT | S | Syn | Para | 25.5 | 17.3 | 0.0092 | 271 | 324 | 40 | 102 | 52 | 41 |
| OUT | T | Para | Allo | 26.8 | 11.0 | 0.0026 | 399 | 218 | 148 | 15 | 62 | 16 |
| OUT | T | Syn | Allo | 26.9 | 11.0 | 0.0009 | 349 | 218 | 80 | 15 | 60 | 16 |
| OUT | T | Syn | Para | 26.9 | 26.8 | 0.74 | 349 | 399 | 80 | 148 | 60 | 62 |

### Simulations

Spatially explicit, individual-based forward simulations (Fig. S1) were performed to help in interpretation of the results. First, we tested whether the observed patterns of MHC similarity between species and between transects could be due to BS alone, without introgression. Second, we checked whether under BS, introgression following secondary contact and hybridization was capable of producing the observed patterns. Third, we explored the temporal and spatial dynamics of introgression under BS and compared them to neutral introgression.

The number of alleles maintained within species depended on the deme size and strength of selection, but the effect of intraspecific migration rate was small, except for the largest deme size (Fig. S5A). Higher migration rate increased the fraction of alleles shared between transects (Fig. S5B). We therefore fixed migration at Nm = 2.5 which, by increasing similarity between transects within species, made our simulations conservative, as stronger introgression would be required for similarity between species within transect to exceed similarity between transects within species. For the same reason we used the initial number of alleles, na = 500, which limited similarity between species at the onset of hybridization, because a large fraction of alleles was lost from each species, independently, during evolution in isolation. Considering allele sharing and AMOVA results simultaneously, balancing selection without hybridization did not produce patterns similar to those observed (Fig. 3). Most importantly, the fraction of alleles shared exclusively between transects within species was always much higher than between species within transect (Fig. 3A). AMOVA results were more complex, as the between transects within species AMOVA component could be smaller or larger than the between species component, depending on the parameter values (Fig. 3B). With increasing deme size and increasing selection both components decreased as more and more variation was distributed within populations.

**Fig. 3.**
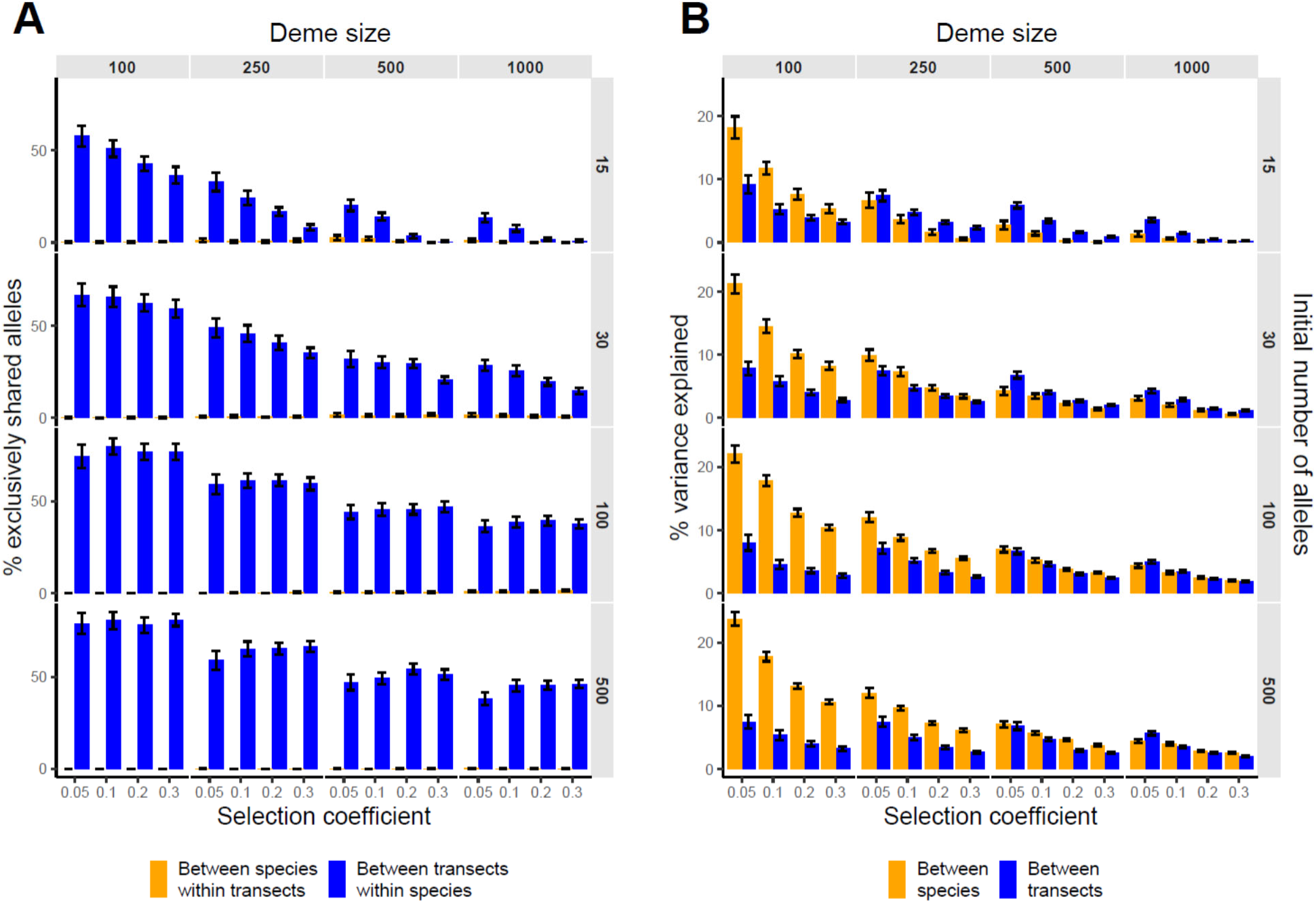
Simulations: Isolation. In all simulations migration rate between adjacent demes within species was Nm = 2.5. A) Percentage of alleles shared exclusively (those shared by two focal groups but absent from all other groups) between species within transect and between transects within species, B) Percentage of total variance explained by between species and between transect within species AMOVA components. Means and 95% confidence intervals from 50 simulations are shown.

Scenarios allowing both BS and hybridization produced patterns similar to the observed under a relatively broad range of parameter values (Fig. 4A, B). The fraction of alleles exclusively shared between species within transects increased with time of hybridization, and after several hundred generations exceeded the fraction of alleles exclusively shared between transects within species (Fig. 4A). Similarly, with increasing time of hybridization, the between transects within species AMOVA component exceeded the between species component, and, as in the observed data, the between species component was negative for the smallest deme size (Fig. 4B). Importantly, introgression of genes under BS was strong even when hybridization was too short or too limited to cause noticeable introgression of neutral genes (Fig. 4C). However, several hundreds of generations were needed for introgression to occur when the hybrid zone constituted a strong barrier (Fig. 4C). Results of simulations assuming multilocus MHC haplotypes were very similar to those for a single multiallelic locus (Fig. S6).

**Fig. 4.**
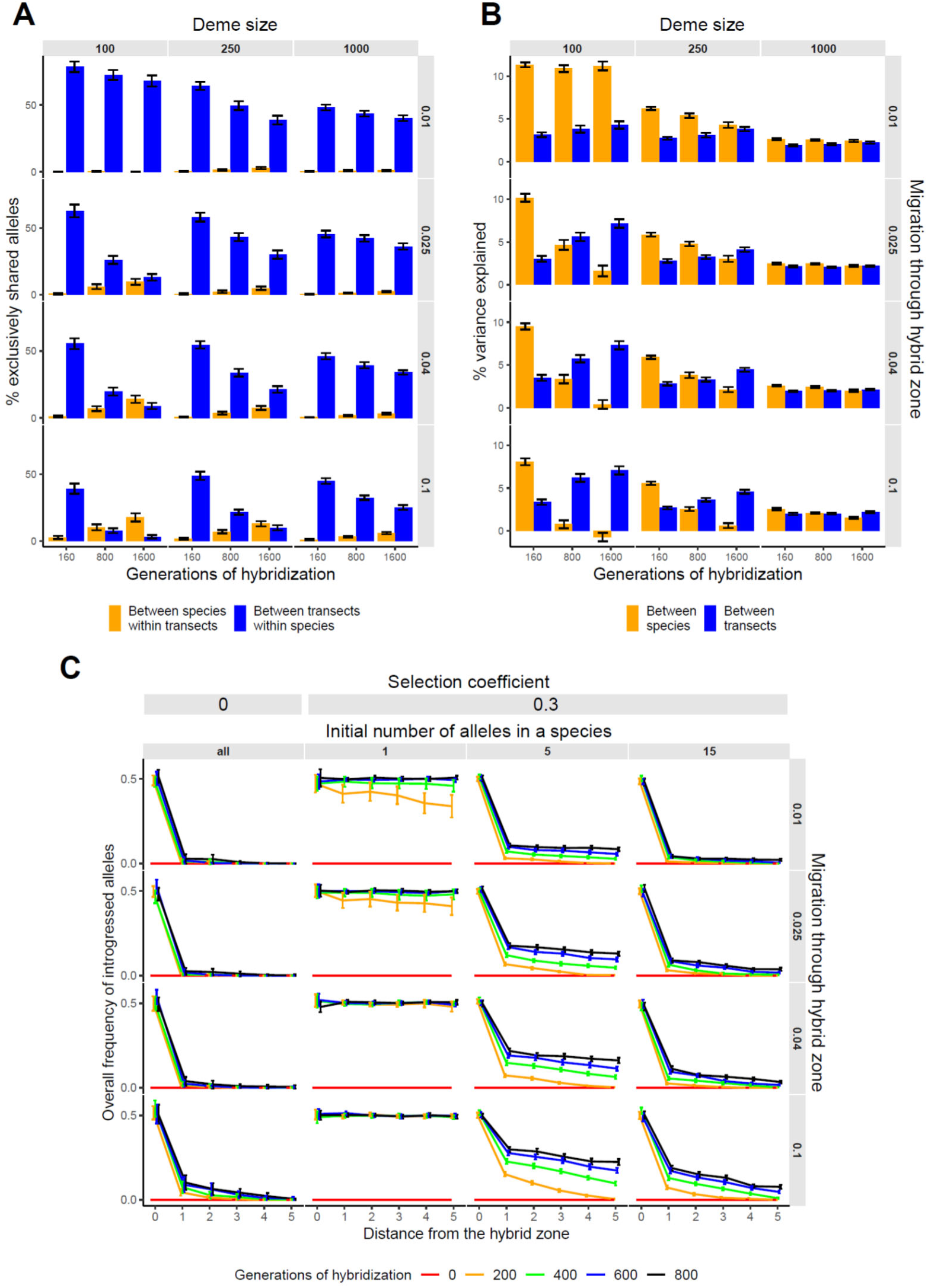
Simulations: Hybridization. In all simulations migration rate between adjacent demes within species was Nm = 2.5. A) Percentage of alleles shared exclusively (those shared by two focal groups but absent from all other groups) between species within transect and between transects within species, B) Percentage of total variance explained by between species and between transect within species AMOVA components. C) Dynamics of introgression in a single transect – frequency of introgressed alleles along the transect at different times after hybridization. As expected, identical results were obtained for neutral introgression, regardless of the initial number of alleles (all). Means and 95% confidence intervals from 50 simulations are shown.

## Discussion

### Evidence for introgression and alternative explanations

This work is, to our knowledge, the first analysis of MHC introgression through hybrid zones. Introgression was detected in both zones, but its extent differed considerably. In the IN zone MHC similarity between species within transects exceeded similarity between transects within species. This implies pervasive introgression: a massive exchange of MHC alleles between species, not limited to specific haplotypes or to a restricted geographic region. The evidence is convincing because the two alternative explanations for the observed pattern – chance and parallel selection – are unlikely. The first was rejected by simulating negative frequency dependence under isolation. The other explanation would require: i) the long-term maintenance of a very large number of alleles across the ranges of both species, and ii) locally similar pathogen communities leading to parallel selection, and parallel changes in allele frequencies Such alleles should, however, be maintained only at low frequencies, otherwise they would be detected as shared between transects within species. This is extremely unlikely, as the number of alleles maintained under BS, although substantial, is limited (Schierup, 1998, our simulations). An additional argument against parallel selection is that there was only a modest increase in local allele sharing in the OUT zone.

### MHC differentiation within transects

Despite pervasive intogression, some MHC differentiation between species was maintained within each transect. The similar location and width of the MHC and the genome-wide clines are remarkable. This apparent similarity may however mask important differences between the two clines. While there is little doubt that the genome-wide cline is secondary, i.e. has been maintained since the formation of the zones after the secondary contact, the origin of the currently observed MHC clines appears more complex. At regional scales, introgression increases both MHC allele sharing and similarity of allele frequencies between species. With limited hybridization, a steep transition between the two MHC pools has probably been maintained since secondary contact, but the nature of this transition may have been evolving. First, the magnitude of differentiation has probably decreased with time since contact, but is still detectable due to limited hybridization and the excellent resolution of highly polymorphic MHC (cf. Fig. 2E, F). Second, at different times, different alleles would contribute to the differentiation. The MHC cline can thus be viewed as both secondary, maintained since contact, but also as primary, being constantly re-formed *de novo* because of a turnover of alleles responsible for the differentiation at any given moment. With time, an increased similarity of allele pools could reduce the selective advantage of introgressed alleles and, in consequence, slow down introgression. This, though, would not affect the above scenario qualitatively.

### Differences between the two hybrid zones

MHC introgression was detected in all transects, but it was more limited in the OUT zone, despite broad syntopy there. The following factors, not mutually exclusive, may differ between the zones and contribute to the observed pattern: i) frequency of hybridization; ii) age of contact; iii) strength of selection favoring MHC introgression; iv) strength of the genomic barrier to MHC introgression.

There could simply be too little hybridization in the OUT zone to allow extensive MHC introgression, even if favored by selection. This is supported by our simulations, and by our empirical data: the presence of admixed individuals, though rare even in syntopy, indicates non-zero recent or ongoing hybridization (Zieliński et al., 2019). Introgression from the *L. vulgaris* lineage inhabiting the OUT zone has been strong enough to cause a regional mtDNA replacement in *L. montandoni* (Zieliński et al., 2013). Furthermore, extensive hybridization was detected in a hybrid zone in southern Poland, where the same *L. vulgaris* lineage hybridizes with *L. montandoni* (Babik et al., 2003). Thus, a negligible frequency of hybridization as the factor limiting introgression is unlikely. Analyses of mtDNA (Pabijan et al., 2017; Zieliński et al., 2013) and nuclear (Pabijan et al., 2017; Zieliński et al., 2016) sequences revealed differences in the history of hybridization between *L. montandoni* and the two *L. vulgaris* lineages. Direct evidence for longer hybridization in the IN zone comes from the observation of shorter genomic heterozygosity tracts in diagnostic markers than in the OUT zone (Zieliński et al., 2019). Within the OUT zone, MHC introgression in the more northern S transect was detectable only in syntopy, while in the more southern T transect also in parapatry. Phylogeographic evidence suggests postglacial colonization of the OUT zone by *L. vulgaris* from the south (Zieliński et al., 2013), so the contact in the S transect could be more recent. However, if the OUT zone originated during postglacial colonization several thousand generations ago, it is difficult to imagine large differences in the age of the contact between the two areas separated by ca. 100 km, and without noticeable barriers to dispersal. The differences in the age of contact in different transects, could, however, be due to anthropogenic habitat alterations during the last thousand years. It is not known how stable the locations of the zones have been, but we have not detected signs of major hybrid zone movement (cf. Wielstra et al., 2017; Zieliński et al., 2019). At first sight longer history of hybridization would make introgression easier to detect. It may, however, not necessarily be so, as alleles derived from older introgression may be lost, or have spread through the species range. The latter would enter the pool of alleles shared between species globally, not locally, and the local sharing was used to infer introgression in the IN zone. Indeed, continent-scale comparisons indicate considerable MHC introgression involving the *L. vulgaris* lineages from both IN and OUT zones (Fijarczyk et al., 2018). Therefore, although old MHC introgression is likely, results from the IN zone support recent and/or ongoing introgression. Differences in the strength of selection or in the distribution of the barrier loci surrounding MHC are also plausible explanations for differences in the extent of introgression between the two hybrid zones. Unfortunately, we do not have enough information to assess the contribution of these factors.

### The nature of selection driving introgression

Pervasive MHC introgression in the face of strong genome-wide isolation implies an adaptive process. There is strong theoretical (Muirhead, 2001) and empirical (Phillips et al., 2018; Pierini & Lenz, 2018) support for novel MHC variants conferring selective advantage. Such advantage may be a major contributor to selection that can overcome strong genome-wide barriers and facilitate adaptive introgression. Adaptive introgression may, in turn, be a major contributor to the complex suite of selective pressures that drive spatial patterns of MHC diversity. For example, simple variant-preserving balancing selection predicts lower-than-neutral structure in the loci of interest, a prediction supported by studies of the S-genes of plants (Glémin et al., 2005). The MHC, in contrast, shows diverse outcomes (reviewed in Spurgin & Richardson, 2010), of which the considerable within-species differentiation by population/region in our study is an example. Also simulations of host-pathogen coevolution showed a rapid turnover of MHC alleles, not consistent with pure negative frequency dependence (Ejsmond & Radwan, 2015). Explicit geographic population structure with some degree of gene exchange may be required to reconcile these properties and processes. Therefore, to obtain a more realistic picture of selection on MHC it is necessary to model MHC-pathogen coevolution in geographically structured populations (Ejsmond, Phillips, Babik, & Radwan, 2018; Ejsmond & Radwan, 2015; Thompson, 2005). Though the pure negative frequency dependence and spatial structure of our simulations were certainly an oversimplification, our settings were likely conservative, and probably made retention of ancestral polymorphism easier and introgression more difficult than in natural systems. We thus conclude that adaptive interspecies introgression may be an important component of the evolution of the MHC.

### Implications

Although the number of studies reporting MHC introgression is still small, several cases described from diverse taxa (Abi-Rached et al., 2011; Grossen et al., 2014; Nadachowska-Brzyska et al., 2012) suggest that the phenomenon may be widespread. Signals of introgression in MHC appears stronger than in other targets of BS (Fijarczyk et al., 2018), though such processes may be more detectable at the MHC. An exciting possibility emerges that MHC may universally be among the last genes to stop flowing between incipient species as reproductive isolation becomes complete, as suggested previously for S genes in plants (Castric, Bechsgaard, Schierup, & Vekemans, 2008). A comparative analysis of multiple vertebrate hybrid zones between strongly isolated but still hybridizing taxa could be used to test this hypothesis. If confirmed, it would support a previously suggested idea (Fijarczyk et al., 2018) of a shared pool of adaptive variation available for hybridizing species that may boost adaptive potential.

Widespread introgression may also contribute to an explanation of why trans-species polymorphism (TSP) seems unusually common in MHC. While TSP has traditionally been attributed to a long-term retention of ancestral polymorphism caused by balancing selection (Klein, Sato, & Nikolaidis, 2007), this explanation is not entirely convincing. Both rapid turnover of MHC alleles observed in empirical surveys and recent simulation results (Ejsmond et al., 2018; Ejsmond & Radwan, 2015), suggest that long term survival of allelic lineages required to explain TSP without introgression may not be as common as previously thought.

## Acknowledgments

We thank S. Bury, D. Cogălniceanu, A. Fijarczyk, M. Liana, M. Niedzicka, M. Pabijan, O. Zinenko for help in sampling. A. Fijarczyk, M. Konczal, M. Pabijan, G. Palomar, K. Phillips, J. Radwan, B. Wielstra and members of our Genomics and Experimental Evolution group provided valuable comments. This work was supported by the Polish National Centre grant 2016/21/N/NZ8/00922 to K.D. and partially supported by the Polish National Centre grant 2014/15/B/NZ8/00250 to P.Z and by the Jagiellonian University (DS/WB/INoS/762/18).

## Data Accessibility

Individual genotypes and sequences of all MHC alleles will be deposited in Dryad upon acceptance.

## Author contributions

K.D, T.S.G. and W.B. designed research, K.D. collected the data, K.D. P.Z., and W.B. analyzed the data, T.S.G. and W.B. designed simulations, T.S.G. performed simulations, W.B. and K.D wrote the paper. All authors read and approved the final version.

