## Supplementary Methods & Figures for "Pervasive introgression of MHC genes in newt hybrid zones"

### *MHC class II genotyping*

MHC class II was amplified in 10 ul PCR reactions containing: 50-100 ng of genomic DNA, 5 ul of Multiplex PCR kit (Qiagen) and each of four primers (IIex2_Fb TCTCTCCRCAGYGGACTTYG, IIex2_Fc TCTSTCCTCAGATGATTTYG, IIex2_R2 CTCACGCCTCCGKTKGTACAGG IIex2_R3 CTCACGCHTCCGSTGCTCCAKG, all sequences 5'->3') at concentration of 1 uM. Individuals were barcoded with a combination of 6 bp indexes at the 5' end of forward and reverse primers. PCR conditions were as follows: initial denaturation at 95 °C for 15 min, followed by 35 cycles: 95 °C for 30 s, 55 °C for 30 s and 72 °C for 70 s and final elongation at 72 °C for 10 min. Amplicons were pooled approximately equimolarly based on gel-band intensity, pools were gel-purified, Illumina adaptors were ligated using NEXTflex PCR-Free DNA Library Prep Kit for Illumina (Bioo Scientific), libraries were quantitated with NEBNext Library Quant Kit for Illumina (NEB) and sequenced on Illumina MiSeq (v3 600 cycles kits). Genotyping was performed using the adjustable clustering method implemented in AmpliSAS (Sebastian, Herdegen, Migalska, & Radwan, 2016). Following clustering of sequence variants within amplicons, we considered only variants with per amplicon frequency > 1.0%. Several variants showing no similarity to MHC class II as well as putative pseudogenes (variants with frameshifts or in-frame stop codons) were excluded from further analyses. To estimate repeatability of genotyping 56 (MHC class I) and 62 (MHC class II) samples were amplified and genotyped twice.

### *MHC class II diversity and tests of selection*

Nadachowska-Brzyska et al. (Nadachowska-Brzyska, Zielinski, Radwan, & Babik, 2012) analysed MHC class II exon 2 variation and tested selection in the whole *Lissotriton vulgaris* species complex. In the present study we analysed a longer fragment and intensively sampled only three out of nine evolutionary lineages within the complex. Therefore we report diversity and results of tests of selection using only sequences obtained in the current study. Divergence between alleles was estimated in MEGA7 (Kumar, Stecher, & Tamura, 2016), by calculating nucleotide (Tamura & Nei), synonymous/nonsynonymous (Nei & Gojobori) and amino acid (Poisson-corrected) distances. To test for the signal of positive selection we compared fit of three codon-based models of evolution (M0, M7 and M8) in PAML (Yang, 2007). To speed up computations we excluded singleton alleles from the analyses. Codons under positive selection (posterior probability (PP) > 0.95) were identified using the Bayes Empirical Bayes Procedure in PAML. Location of positively selected codons was compared to that of the ABS in human MHC class II (Tong et al., 2006).

### *Classical MHC alleles and supertypes*

MHC class I alleles were previously classified, on the basis of expression level and sequence similarity, into two classes: i) HEX – intermediate and high expression, putative functional alleles, ii) LEX – low expression, putative nonclassical/nonfunctional alleles (Fijarczyk, Dudek, Niedzicka, & Babik, 2018). New MHC class I alleles detected in the current study were incorporated into this classification using the rules described in (Fijarczyk et al., 2018). For MHC class II information on the expression status is more limited, although most alleles amplified with our primers appear to be expressed (Nadachowska-Brzyska et al., 2012). We have therefore not attempted to classify the class II alleles into putative classical and nonclassical/nonfunctional groups.

The idea of supertype analysis is to cluster alleles into classes of functionally similar sequences as defined by physico-chemical properties of amino acids in positions that determine specificity of antigen binding. Despite an overwhelming signal of positive selection in both MHC classes, only a few codons were identified as positively selected (5 in class I and 2 in class II), too few to define supertypes. Therefore we used the combination of those positively selected codons and previously described human ABS (Reche & Reinherz, 2003; Tong et al., 2006). Five physicochemical descriptors of each amino acid (Sandberg, Eriksson, Jonsson, Sjöström, & Wold, 1998) were used to group alleles using the K-means clustering in *adegenet* (Jombart, 2008). Based on the Bayesian Information Criterion (BIC) we identified 25 supertypes for class I (only HEX alleles were included) and 22 supertypes for class II. We note two limitations of the supertype approach as applied to the newt dataset. First, because of the small number of codons identified as positively selected, supertypes were based mostly on human ABS positions. Second, the value of BIC decreased monotonically with the increasing number of supertypes, making identification of the number of supertypes somewhat arbitrary. The supertypes should thus be regarded as clusters of alleles based on amino acid similarity in most polymorphic positions rather than robustly defined groups of functionally similar alleles. As a consequence, supertype-based results should be treated with caution.

### *Simulations*

Spatially explicit, forward in time simulations were performed using Selector (Currat, Gerbault, Di, Nunes, & Sanchez-Mazas, 2015). Each species was represented by 15 demes exchanging migrants according to the stepping stone model, i.e. only between adjacent demes. Two transects, 5 demes each, were connected by 5 vertically arranged demes (Fig. S1). Because the distance between transects in the IN zone was approximately 5 x larger than the length of the transects, to reduce computational burden, migration between the connecting demes was set to ca. 0.27 of that between demes within transects (Charlesworth & Charlesworth, 2010, eq. 7.8a), so that the 5 connecting demes actually corresponded to 25 demes exchanging migrants at the same rate as demes within transects. Three strengths of migration between demes within transect were evaluated (Nm = 0.1, 0.5, 2.5).

At the beginning of the simulations (Fig. S1I), only a single, centrally located deme within each species was occupied (N0 = 1000 individuals) and both species shared a single pool of alleles (na = 15 – 500). Selector starts with a uniform allele frequency distribution, so the initial differences between the species were minor, resulting only from sampling error. This setting emulated the split of a single ancestral species into two descendant species of equal sizes. Note however that the number of alleles maintained within each species was limited (Fig. S5), so depending on the initial number of alleles, large fraction of alleles could be lost from each species, quickly reducing the number of shared alleles. No mutations were allowed throughout simulations. Various strengths of negative frequency dependent selection (s = 0.05 – 0.3) were evaluated and selection was kept constant throughout a simulation; fitness of an allele with frequency $f(a)$ was defined as $1 – f(a)s$. Following establishment of the founding populations, colonization of the initially empty demes (identical carrying capacities, N = 100 – 1000, growth rate 0.5) within each species and migration between demes occurred for 1000 generations. Then the carrying capacity of the founding demes was set to that of other demes (Fig. S1II). Both species were evolving in isolation for further 15 000 generations and then hybridization was allowed by setting carrying capacity of two, previously unoccupied, demes located in the centre of each transect to 0.01 – 0.1N (Fig. S1III). Thus, in each transect the hybrid zone consisted of a single deme. Immigration into the zone was high and symmetrical, i.e. demes adjacent to the zone sent identical number of emigrants in each direction, and emigration from the zone was reduced to 0.01 – 0.1 of that value (controlled by the carrying capacity of the hybrid zone population), making the zone a barrier to introgression. Hybridization was allowed for 160 - 1600 generations corresponding to 0.01-0.1 of the time of evolution in isolation. For each combination of parameter values 50 simulations were performed.

At the end of each simulation samples of 16 individuals were taken from each deme within transect (except of the hybrid zone deme, Fig. S1IV) and the following statistics were calculated: i) percentage of variation explained by the between species and between transects within species AMOVA components, ii) fraction and number of alleles shared between species and between transects within species, iii) fraction and number of alleles shared exclusively (those shared by two focal groups but absent from all other groups) between transects within species.

To investigate the temporal and spatial dynamics of introgression we recorded the fraction of introgressed gene copies in each deme within the transect at various times following hybridization. A single transect with demes of N = 250 and distinct (non-overlapping) sets of alleles in each species was simulated, which allowed straightforward calculation of the fraction of introgressed gene copies. Scenarios with a single allele initially fixed within each species and with 5 and 15 alleles per species were investigated. The latter reflected the number of alleles maintained within species for deme size N = 250 individuals under strong negative frequency dependent selection (Fig. S5). Immediately after establishment of populations we turned on hybridization and recorded, using actual allele frequencies reported by Selector, the fraction of introgressed gene copies in each deme at different times following the onset of hybridization. We investigated both neutral and negative frequency dependent scenarios, s = 0 and 0.3, respectively and different strengths of introgression, 0.01-0.1 of migration between demes within species ($N_hm$ = 0.025-0.25). For each combination of parameters 50 simulations were performed.

Simulations described above assumed a single multiallelic locus, but MHC in newts is multilocus and both classes are tightly linked. To check for the effect of these differences in genetic architecture between simulated and real data, we ran a subset of scenarios modelling multilocus MHC haplotypes explicitly. The following parameter values were investigated: s = 0.3, Nm = 2.5, na = 500 (in this case the initial number of multilocus haplotypes), N= 100, 250, 1000, $N_hm$ = 0, 0.04, 0.1 and the time of hybridization was set to 0.1 of the time of evolution in isolation.

To obtain haplotypes for simulations we used the following procedure. First, the empirical distributions of (i) the number of MHC alleles per individual (class I and II combined) and (ii) allele frequencies, were constructed with both species combined, but excluding mixed populations. Second, we estimated parameters of (iii) such a normal distribution describing the number of MHC alleles per haplotype, that produced, when alleles on haplotypes were sampled from (ii), the distribution of the number of alleles per individual most similar to (i). At the beginning of each Selector simulation, a haplotype with the number of MHC alleles sampled from distribution (iii) and their identity sampled from the allele frequency distribution (ii) was assigned to each allele simulated by Selector. Following the simulation AMOVA was performed with the number of pairwise differences between haplotypes as the distance measure.

Supplementary Tables S1 to S4 are in a separate Excel Workbook

**Table S1. Sampling and MHC variation.**
**Table S2. Sequence divergence between MHC class II alleles.**
**Table S3. AMOVA for supertypes.**
**Table S4. MHC and genome-wide cline parameters.**

**Supplementary Figures**

Fig. S1. **Design of simulations.** Four stages of simulations shown in panels I – IV were: I) an ancestral species split into two equally sized, completely isolated descendant species, which thus initially shared a pool of alleles, II) following the split, each species colonized a world of demes conforming to a one-dimensional stepping stone model arranged into a horseshoe shape that approximated the two transects of the IN zone, i.e. the distance between transects was 5 times larger than the length of a transect within each species, III) following prolonged evolution in isolation, secondary contact and hybridization ensued; a single deme hybrid zone acted as a partial barrier to gene flow – immigration into the zone was high but emigration from the zone was strongly reduced as in classical tension zone models. Details of simulations are in Supplementary Methods.

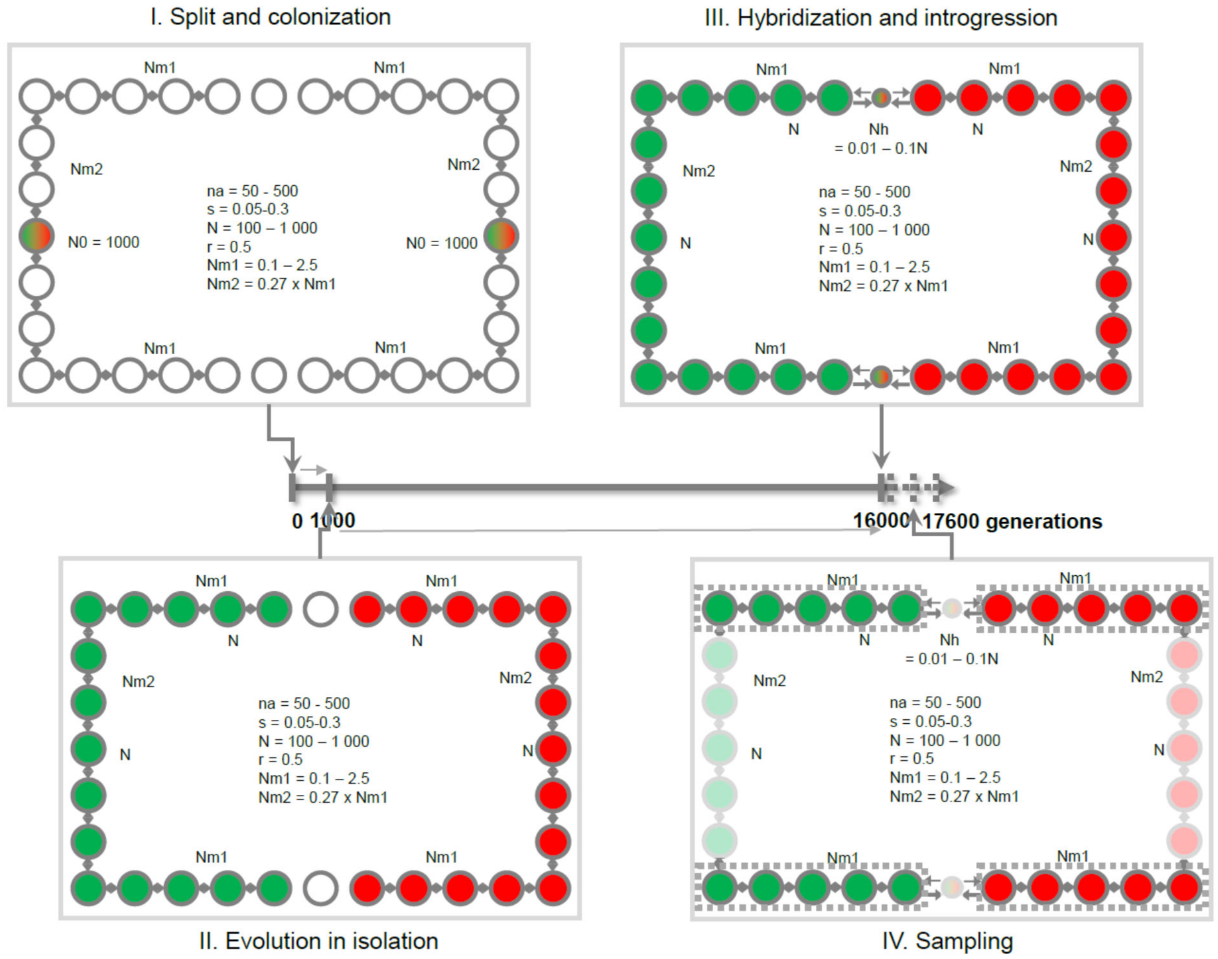

Fig. S2. **Per individual number of MHC alleles**. Red - *L. montandoni*, green – *L. vulgaris*; Individuals from syntopic populations were excluded. Dashed lines show the averages. A) MHC class I, B) MHC class II.

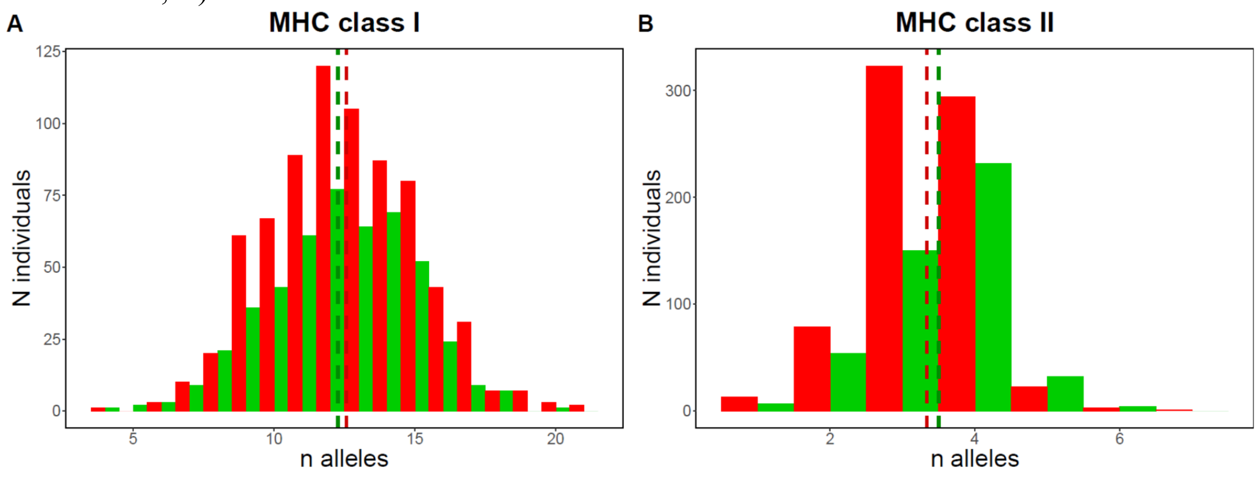

Fig. S3. **Sequence logo summarizing amino acid variation among sequences of MHC class II alleles.** Positions identified as the Antigen Binding Sites (ABS) in all MHC class IIB genes are highlighted in yellow. Asterisks denote codons under positive selection. Note high diversity in most ABS positions.

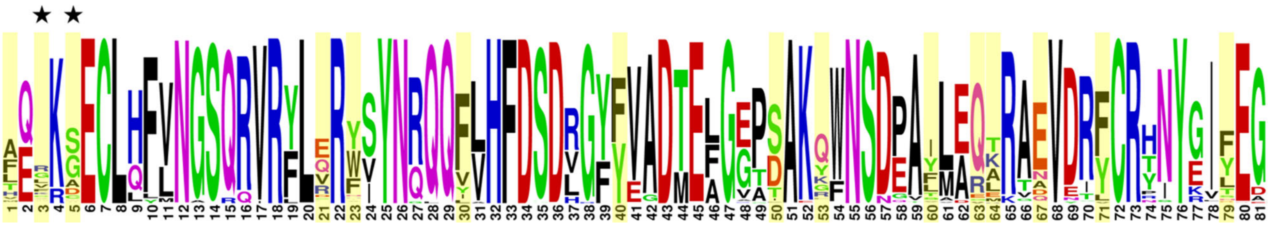

Fig. S4. **Principal Component ordination of individuals based on MHC data**. In all plots individuals from syntopic populations were excluded; light green – *L. vulgaris* inside the Carpathian Basin (IN), dark green – *L. vulgaris* outside the Carpathian Basin (OUT), light red – *L. montandoni* IN, dark red – *L. montandoni* OUT. PCAs are based on: A) all MHC class I alleles, B) all MHC class II alleles, C) Putative functional class I alleles (HEX), D) putative nonclassical/nonfunctional MHC class I alleles (LEX).

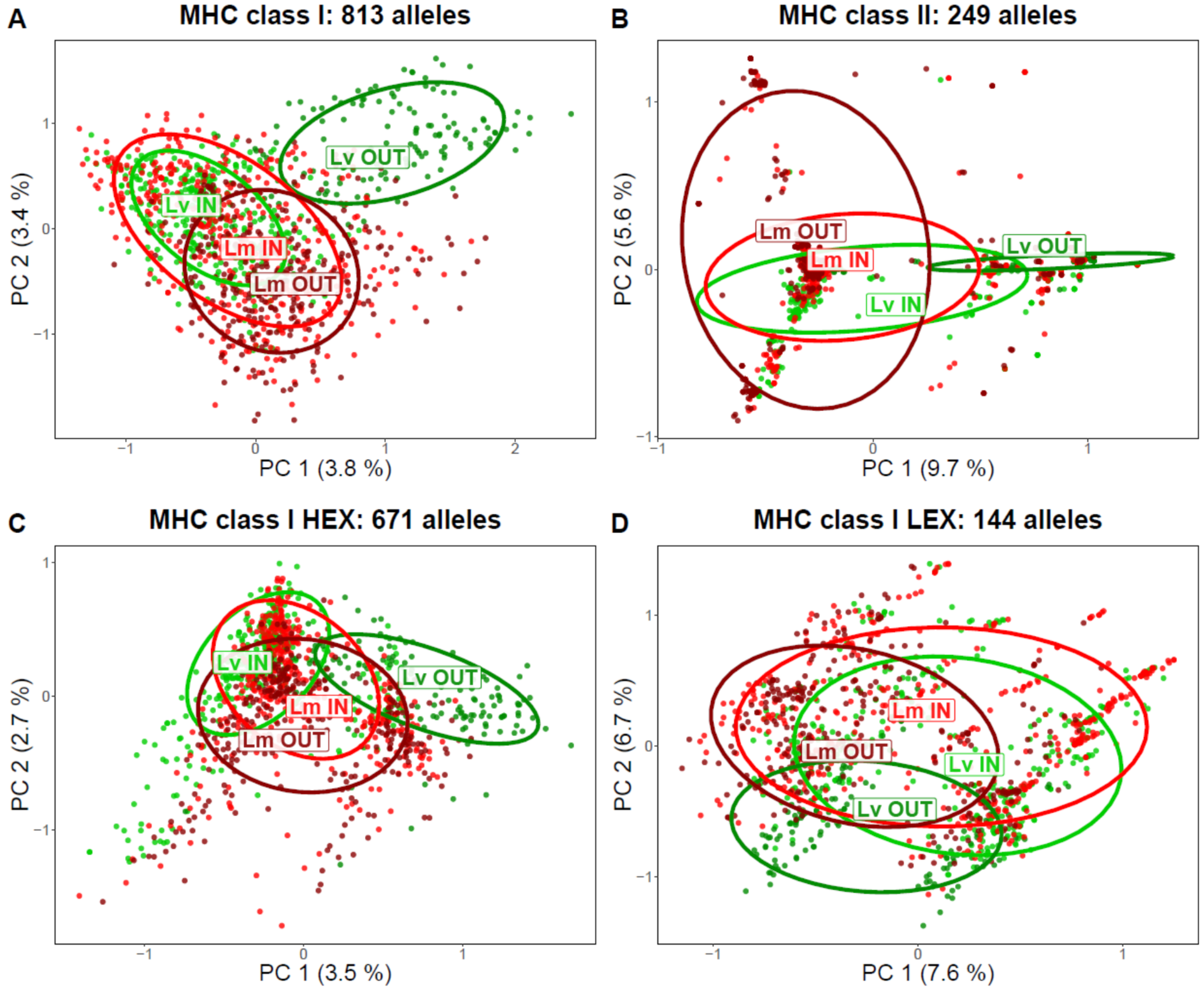

Fig. S5. **Relationship between MHC variation and intraspecific migration.** All results for scenarios without hybridization, initial number of alleles na = 500, after 16 thousand generations; migration rate expressed as a product of deme size and migration rate (Nm). A) Number of alleles maintained within species, B) Number of alleles shared between transects within species. Means and 95% confidence intervals from 50 simulations are shown.

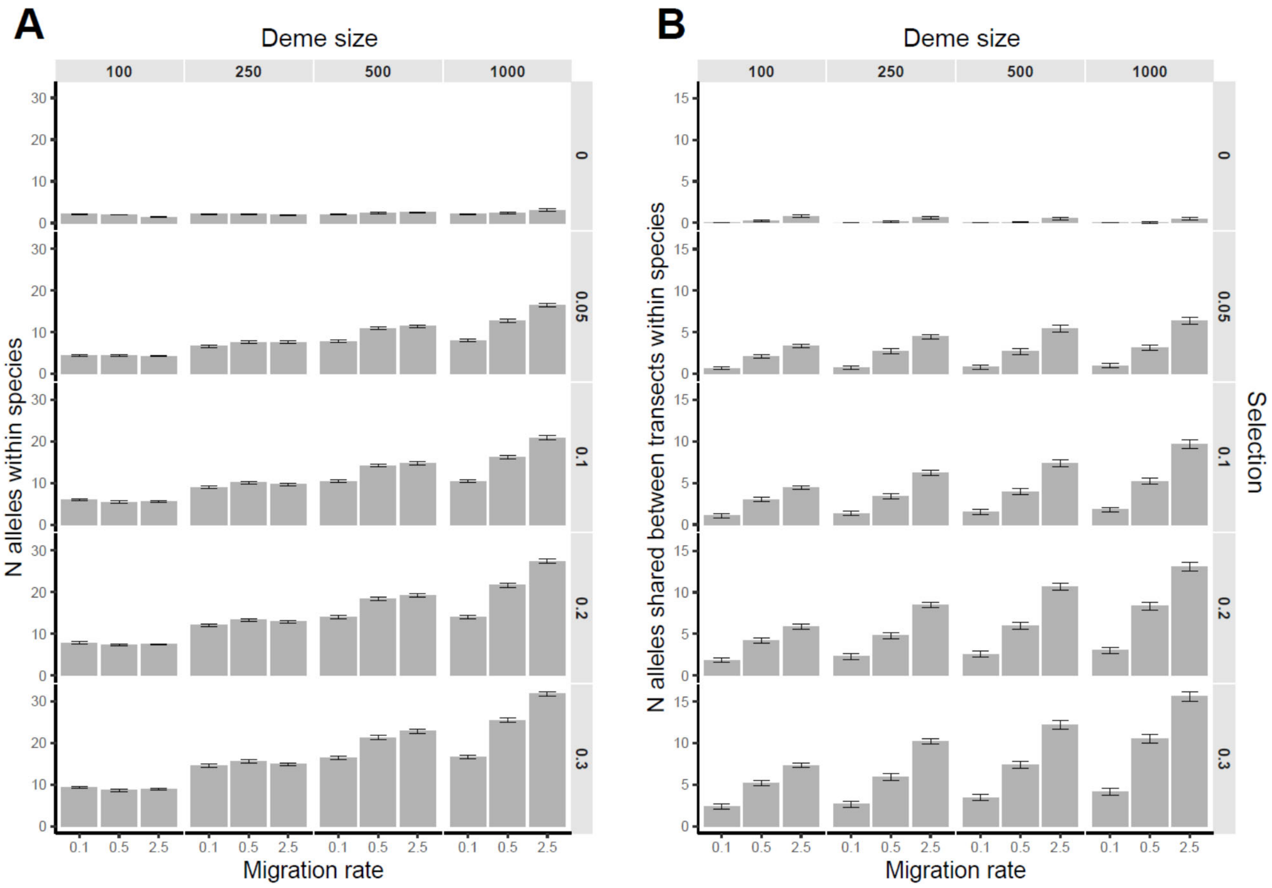

Fig S6. **Comparison of simulation results with MHC modelled as a single multiallic locus (A, C) or as multilocus haplotypes (B, D).** For each haplotype the number of alleles was sampled from the normal distribution with mean = 7.6 and variance = 3.04, while their identity was sampled from the empirical distribution of allele frequencies (for details see Supplementary Methods).A, B: Percentage of total variance explained by between species and between transect within species AMOVA components. C, D: Percentage of alleles shared exclusively (those shared by two focal groups but absent from all other groups) between species within transect and between transects within species.

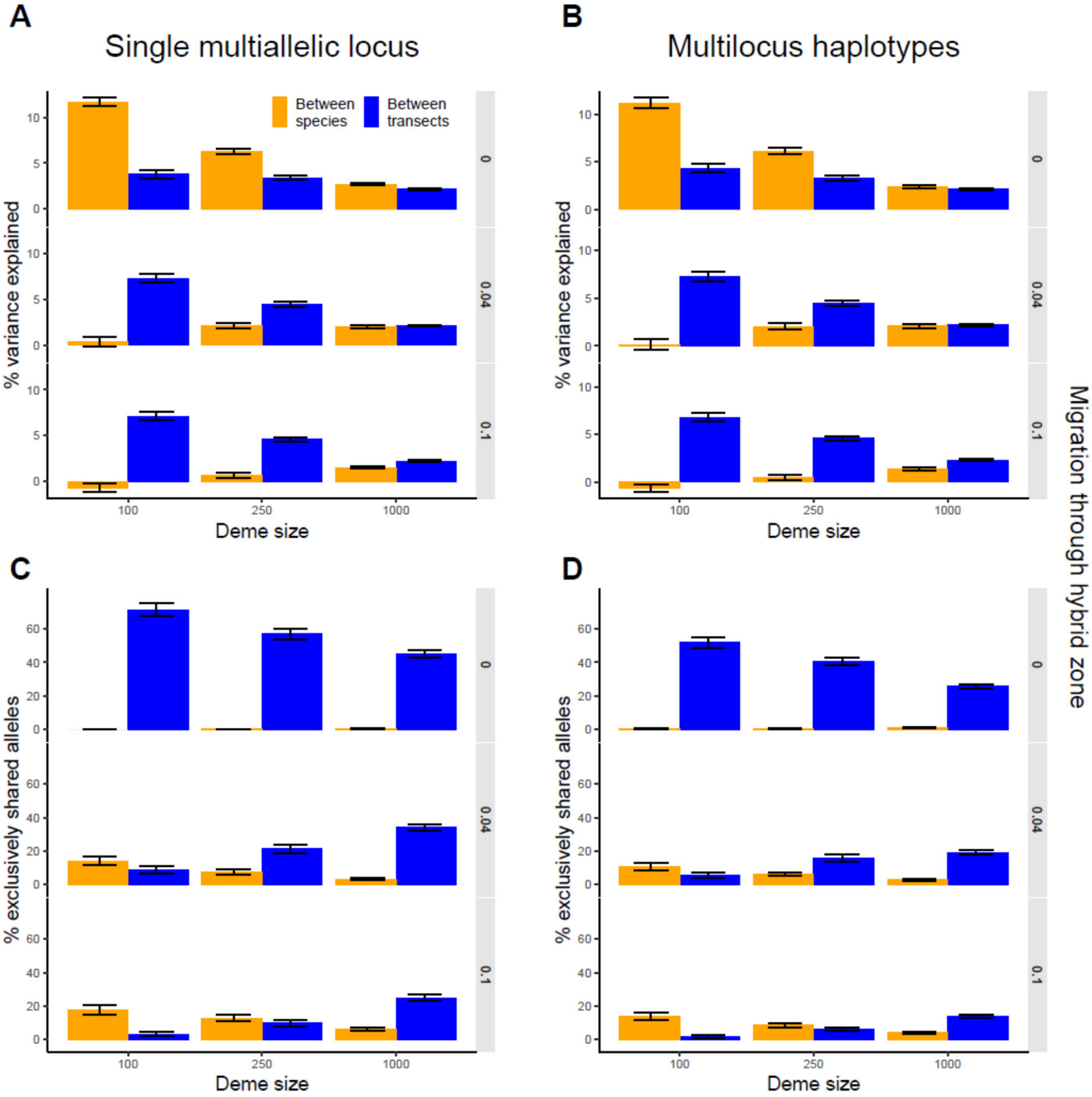
